# Hidden species diversity in a living fossil vertebrate

**DOI:** 10.1101/2022.07.25.500718

**Authors:** Chase D. Brownstein, Daemin Kim, Oliver D. Orr, Gabriela M. Hogue, Bryn H. Tracy, M. Worth Pugh, Randal Singer, Chelsea Myles-McBurney, Jon Michael Mollish, Jeffrey W. Simmons, Solomon R. David, Gregory Watkins-Colwell, Eva A. Hoffman, Thomas J. Near

## Abstract

Ancient, species-poor lineages persistently occur across the Tree of Life. These evolutionarily unique lineages are likely to contain unrecognized species diversity masked by the low rates of morphological evolution that characterize living fossils [1, 2]. Halecomorphi is a major clade of ray-finned fishes that diverged from its closest relatives over 200 million years ago [3, 4] yet is represented by only one recognized living species in eastern North America, the Bowfin *Amia calva* Linnaeus. Here, we use double digest restriction-site associated DNA (ddRAD) sequencing and high-resolution computed tomography to illuminate recent speciation in the bowfins. Our results support the resurrection of a second living species of Bowfin with the timing of diversification dating to the Pleistocene. In turn, we expand the species diversity of an ancient lineage that is integral to studies of vertebrate genomics and development [2, 3, 5], yet is facing growing conservation threats driven by the caviar fishery [6].

## Main Text

The Bowfin *Amia calva* is the sole living representative of Halecomorphi, an extremely old lineage of ray-finned fishes with a cosmopolitan distribution in the fossil record classically labeled as living fossils [3]. Together with the seven living species of gars, the Bowfin forms the sister lineage to Teleostei, which comprises nearly half of all living vertebrate species [4, 5].

Together with sturgeons, the Paddlefish, and mooneyes, Bowfin and gars form a hotspot of ancient freshwater vertebrate diversity in North America [3, 4].

Because of its evolutionary history, the Bowfin is exceptionally important for understanding genomic, developmental, and immunological evolution in vertebrates [2, 5]. The Bowfin is also notable for its apparently low rates of molecular evolution [2, 5] and phenotypic similarity to extinct species from over 145 million years ago [3]. In addition, the economic significance of Bowfin is rapidly increasing with an intensifying demand for sources of caviar [6], putting pressure on extant populations already strained by the centuries-long reputation of *Amia* as a “rough fish” [7]. Nonetheless, there has been no comprehensive investigation of the genetic or morphological diversity among populations of Bowfin.

To investigate the possibility that the Bowfin includes hidden species diversity, we collected genome-wide double digest restriction-site associated DNA (ddRAD) DNA sequences for 177 specimens and phenotypic data from 255 specimens of *Amia* sampled across the entire present-day distribution of the species (Fig. 1a). We aligned the ddRAD loci using the recently published *Amia calva* genome [5], harvesting a total of 56,247 loci. Our phylogenomic analysis (Fig. 1b) unambiguously resolves two major lineages in *Amia*; one includes specimens from the type locality of *A. calva* in Charleston, South Carolina, USA and the other corresponds to a lineage for which the oldest available name is *Amia ocellicauda* Richardson 1836 (Supplement) from Lake Huron in Ontario, Canada. Estimates of genomic ancestry corroborate the resolution of sampled individuals into two phylogenetic lineages (Fig. 1b) The geographic distribution of the two species of *Amia* is suggestive of allopatric speciation associated with rivers draining into the Gulf of Mexico (Fig. 1a). Pairwise F_ST_ values show much higher genetic differentiation (∼0.35-0.75) between *Amia calva* and *A. ocellicauda* than in comparisons within either species (Fig. 1c). Principal components analysis of single nucleotide polymorphisms harvested from the ddRAD loci between the two Bowfin species also corroborates the presence of deep genomic divergence between the two distinct species lineages (Supplement). Relaxed molecular clock analyses of holostean fishes estimates that the two species of *Amia* diverged during the Pleistocene 1.82 Ma (95% CI: 0.95 to 2.93 Ma; Supplement), which is consistent with a pattern of glaciation-induced speciation in other North American freshwater vertebrates [8].

**Figure 1.**
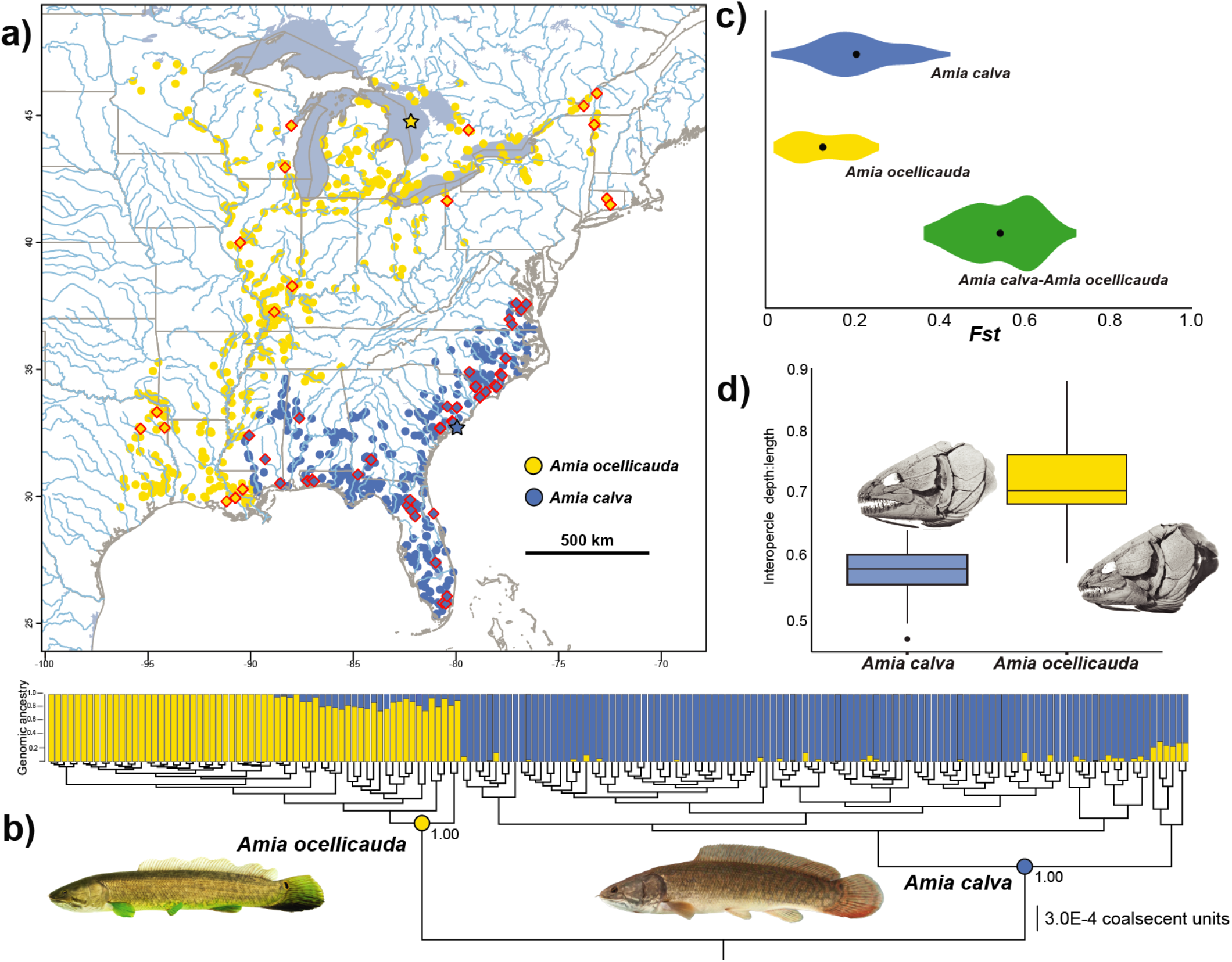
Identification of a hidden Bowfin species diversity. (a) Map of eastern North America showing occurrences of *Amia calva* (blue) and *A. ocellicauda* (yellow). Stars indicate type localities. Diamonds indicate specimens sampled in our ddRAD phylogenetic analysis. (b) Phylogeny and genomic structure analysis of 177 specimens of *Amia* based on 56,247 ddRAD loci, showing genomic ancestry and reciprocal monophyly of both *Amia calva* and *A. ocellicauda*. Photograph of *Amia ocellicauda* from lower Tennessee River, Marshall Co., Alabama USA YPM 035200 by JMM and *Amia calva* from the Suwanee River, Gilchrist Co., Florida USA UF 238466 by Zachary Randall. (c) Comparison of pairwise *F*_*st*_ values for comparisons within each species and the comparisons between *Amia calva* and *A. ocellicauda*. (d) Boxplot showing interopercle robusticity (ratio between maximum dorsoventral depth and maximum anteroposterior length) in *Amia calva* and *A. ocellicauda*. The long lower tail on the *A. ocellicauda* boxplot is created by the inclusion of a juvenile specimen; ontogenetically mature individuals do not overlap. CT-scanned skull of *Amia calva* is TU 22613; CT-scanned skull of *A. ocellicauda* is TU 118772.

A comparison of scale row and fin element counts for 255 specimens of *Amia* found no consistent differences between *A. calva* and *A. ocellicauda* (Supplement). We measured cranial bone proportions in 35 specimens of *Amia*, 18 of which were scanned using high-resolution computed tomography. Specimens of *Amia ocellicauda* consistently possess more robust interopercles than *A. calva* (Fig. 1d) and have 15 dentary teeth rather than 16 or 17 as in *A. calva*. The discovery of two phenotypic traits consistent with the phylogenomic results amplifies our delimitation of two living species of *Amia*.

Our study reveals the presence of two recently diverged sibling species of bowfins. Despite the extremely old age of their parent clade, *Amia calva* and *A. ocellicauda* have diverged in the last two million years, contrasting with the view of the Bowfin as an evolutionary ‘dead-end’ and recalling other ancient lineages that have more recently produced their standing species diversity [9]. A more accurate understanding of species diversity of bowfins will inform conservation decisions for this evolutionarily unique lineage that is the target of an emerging caviar fishery [6]. In turn, the illumination of hidden living diversity in bowfins demonstrates that North America has acted as both a cradle and refugium of ancient vertebrate diversity [3-5]. Along with evidence for deep splits in living lineages of classic living fossils like coelacanths [10], the resurrection of *Amia ocellicauda* reveals the potential for hidden species richness awaiting discovery in other deeply divergent and species-depauperate vertebrate lineages.

## Supplement

Detailed methods and further discussion of the results are available in the Supplement.

## Acknowledgments

This work was supported by the Bingham Oceanographic Fund maintained by the Peabody Museum, Yale University. CT scanning support was provided by the Yale Chemical and Biophysical Instrumentation Center, Eric Paulson, and Brandon Mercado (Yale University). This study utilized tissue holdings curated and maintained at the North Carolina Museum of Natural Sciences and the Yale Peabody Museum. Additional tissue specimens were provided by Jeffrey Stein and Sarah King (Illinois Natural History Survey, University of Illinois), Brady Porter (Duquesne University), W. Leo Smith and Andrew Bentley (University of Kansas), Larry Page and Rob Robins (University of Florida), Matt Wagner (Mississippi Museum of Natural History), Eric Hilton (Virginia Institute of Marine Science), Preston Chrisman (South Carolina Department of Natural Resources), and the Royal Ontario Museum. Specimens and field support were provided by Joseph Ballenger and John Archambault (South Carolina Department of Natural Resources), Mike Beauchene, Edward Machowski, Christopher McDowell, and Lillian Glynos (Connecticut Department of Environmental Protection), Alice E. Near, Rebecca J. Near, and Lake Champlain International.

## Supplemental Inventory Supplemental Figures and Tables

### Supplemental Experimental Procedures

**Figure S1.**
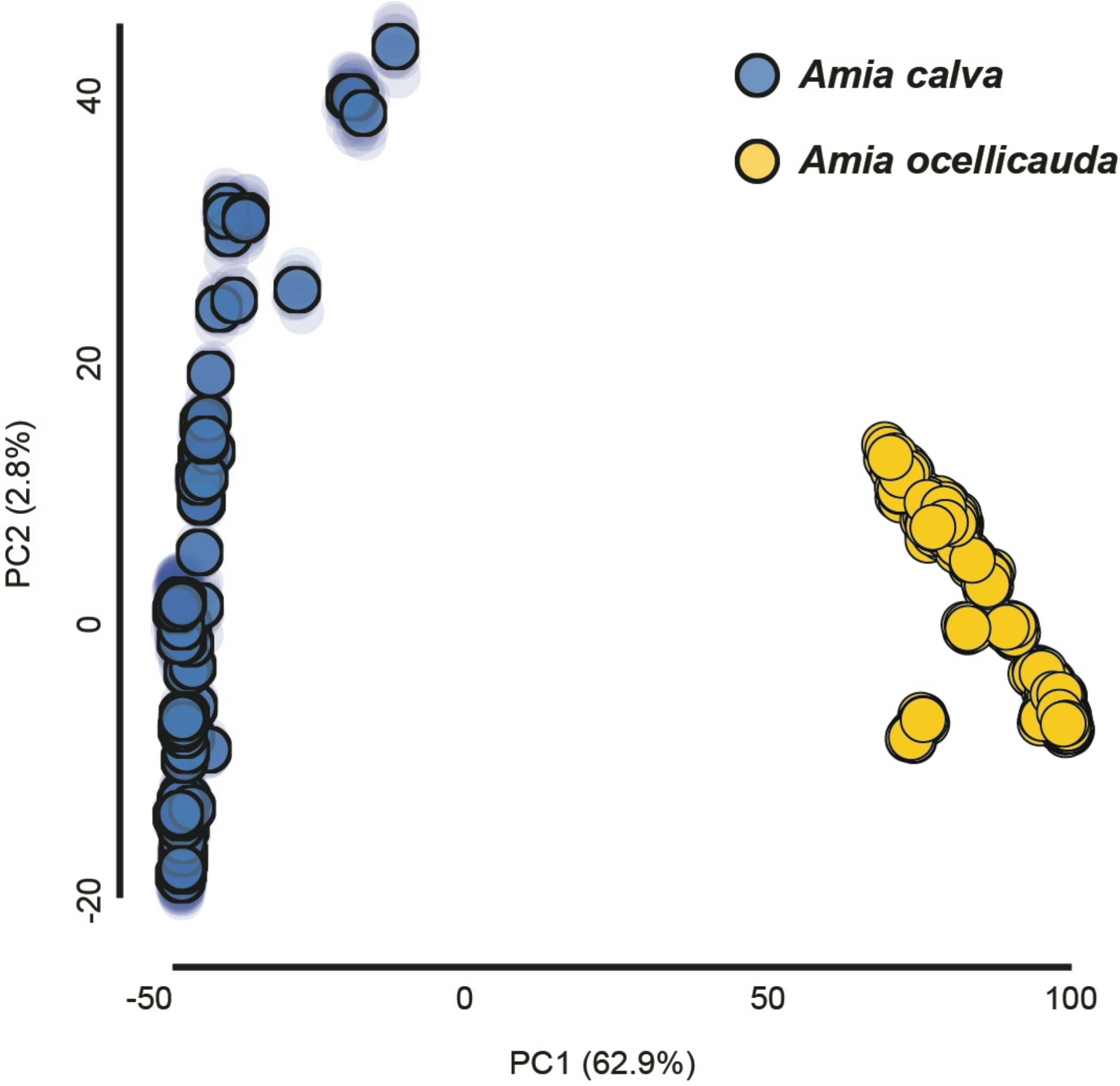
Plot of first and second principal components (PC) of ddRAD loci among the 177 genotyped specimens of *Amia calva* and *A. ocellicauda*.

**Figure S2.**
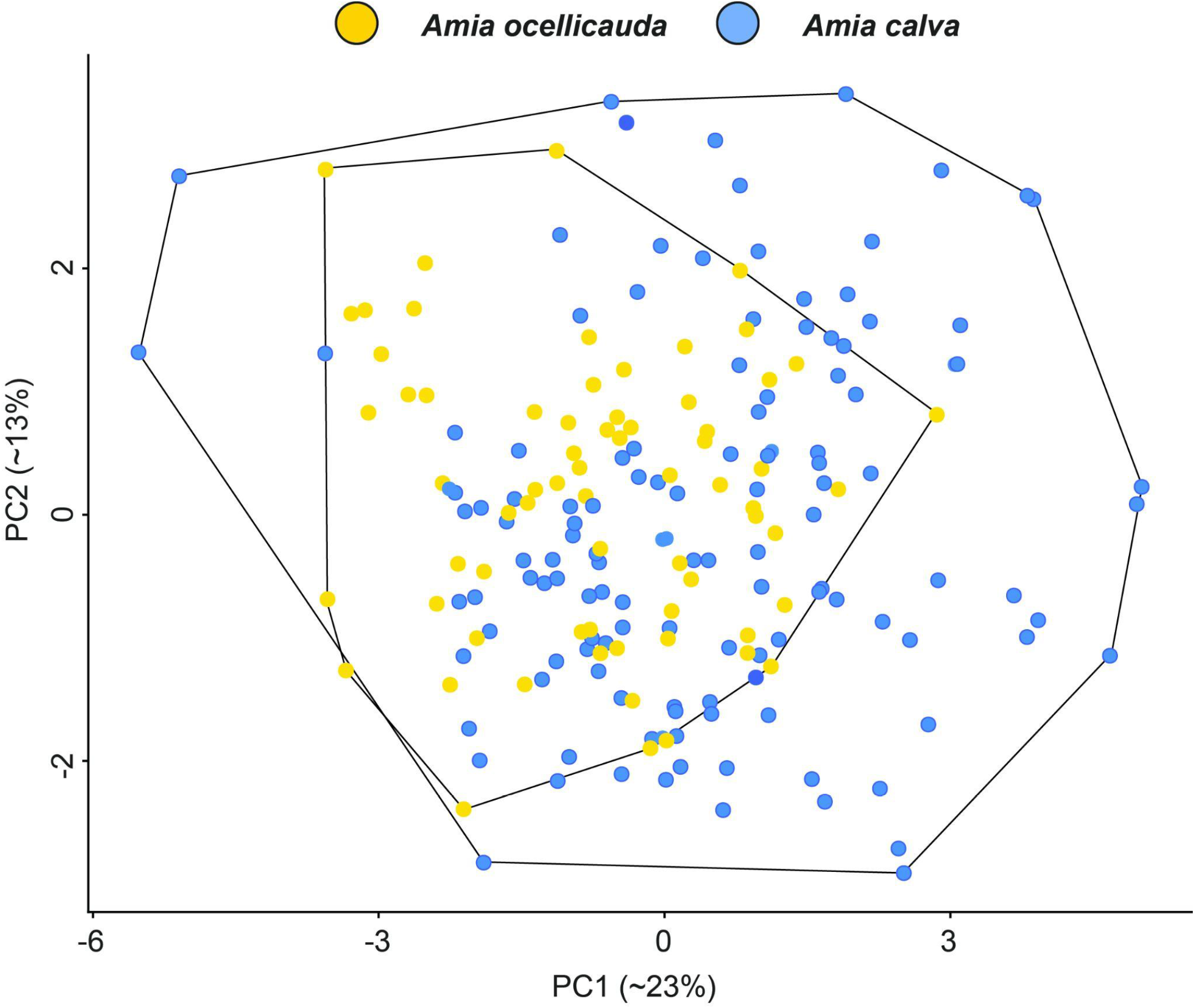
Plot of first and second principal components (PC) of meristic traits for *Amia calva* and *A. ocellicauda*.

**Table S1.** Parameters for specimens scanned using high-resolution computed tomography.

| **Specimen** | **Voltage (kV)** | **Current (uA)** | **Exposure (fps)** | **Voxel size (mm)** |
| --- | --- | --- | --- | --- |
| YPM ICH 27420 | 69 | 66 | 1 | 0.02955711 |
| YPM ICH 34789 | 71 | 67 | 1 | 0.06428329 |
| YPM ICH 34799 | 73 | 71 | 1 | 0.05983228 |
| YPM ICH 34809 | 78 | 68 | 1 | 0.05313689 |
| YPM ICH 35169 | 90 | 86 | 1 | 0.06273028 |
| YPM ICH 35181 | 108 | 107 | 2 | 0.05838511 |
| YPM ICH 35182 | 83 | 79 | 1.41 | 0.0513878 |
| YPM ICH 35189 | 98 | 95 | 1.41 | 0.06971625 |
| TU 118772 | 113 | 111 | 2 | 0.07338808 |
| TU 22613 | 82 | 81 | 1.41 | 0.05215674 |
| TU 28482 | 95 | 96 | 2 | 0.05424211 |
| UF 55734 | 91 | 90 | 2 | 0.05694684 |
| YPM ICH 34815 | 95 | 90 | 2 | 0.05578556 |
| YPM ICH 34816 | 95 | 90 | 2 | 0.05578556 |
| YPM ICH 35173 | 95 | 90 | 2 | 0.05578556 |
| YPM ICH 35184 | 95 | 90 | 2 | 0.07054719 |
| UAIC 03044.01 | 86 | 90 | 2 | 0.06954194 |
| UAIC 04059.07 | 90 | 90 | 2 | 0.05494159 |

### Specimen sampling

Most specimens of *Amia calva* and *A. ocellicauda* were sampled over the course of several field seasons in aquatic habitats using a combination of backpack and boat-mounted electroshockers, and seine nets. Tissue samples were stored in 99% ethanol. Morphological voucher specimens were euthanized, fixed in an aqueous solution of formaldehyde for up to 21 days, soaked in tap water for up to seven days, and transferred to 70% ethanol for long term preservation in the ichthyology collection at the Yale Peabody Museum. Tissue samples were also obtained from museum collections. The sampling locations and museum collection records (if applicable) of all specimens used in the phylogenomic analyses are available on Dryad at http://dx.doi.orgXXXXX.

### Generation of double digest restriction-site associated DNA (ddRAD) loci

We extracted genomic DNA using a Qiagen DNeasy Tissue Extraction Kit (Qiagen, Valencia, CA, USA), following the manufacturer’s protocols. The preparation of ddRAD libraries followed protocols outlined in Peterson et al. [S11]. Approximately 400 ng of each starting DNA sample was digested using PstI/MspI restriction enzymes at 37C for 16 hours, ligated with 96 unique barcodes per plate (96-well) at 22C for 180 minutes, and then amplified using PCR. The PCR amplification conditions consisted of an initial denaturation at 98C for 30 seconds, followed by 16 cycles with denaturation at 98C for 30 seconds, annealing at 62C for 30 seconds, and elongation at 72C for 30 seconds, and a final elongation at 72C for 10 minutes. We normalized DNA concentrations across samples after ligation and PCR amplification steps for all samples in each plate, purified using a QIAquick PCR Purification Kit (Qiagen, Valencia, CA, USA), and pooled before proceeding to the next step. Purified PCR products for each batch were pooled into one library and size-selected for fragments ranging between 300–500 bp using a BluePippin 2% cassette (Sage Science, Beverly, MA, USA). Each size-selected library was validated using a 2100 Bioanalyzer (Agilent, Santa Clara, CA, USA) and DNA fragments were sequenced using 100 bp single-end sequencing on an Illumina NovaSeq 6000 at the University of Oregon GC3F facility (http://gc3f.uoregon.edu). The demultiplexed reads were run through ipyrad v.0.9.68 [S12], using default setting with the following exceptions: ‘denovo’ for assembly method, ‘ddrad’ for datatype, ‘TGCAG, CCG’ for restriction overhang, ‘0.90’ for clustering threshold, and ‘2’ for a stricter adapter filtration. The minimum number of specimens sharing a locus, hereafter referred to as ‘min’, was set to smallest numbers (184 and 75, see below) to reach a stationarity of loci dropout rate. The raw ddRAD sequencing files are available on Dryad at http://dx.doi.orgXXXXX.

### Phylogenomic and population genomic analyses

The phylogenetic relationships among 177 sampled specimens of *Amia* were inferred from a concatenated DNA sequence dataset of the ddRAD loci. A posterior set of relative time calibrated phylogenetic trees were generated using BEAST 2.6.4 [S13] with a coalescent constant population size branching model, a GTR molecular evolutionary model with gamma distribution of among site rate variation, and a strict molecular clock model with a clock rate of 1.0. The Markov chain Monte Carlo (MCMC) was run for 1.0 × 108 generations and log and tree files were updated every 1.0 × 104 generations. Convergence of parameters values in the BEAST 2.6.4 MCMC were assessed by the effective sample sizes that were calculated using Tracer version 1.7. Generations sampled before convergence was attained were discarded as burn-in. The BEAST analyses were run three separate times and post burn-in generations were pooled from all three runs using LogCombiner 2.8. A maximum clade credibility tree with median node heights was constructed for the post burn-in species tree topologies using TreeAnnotator 2.6.4. The xml files used in BEAST analyses and the summarized tree file are available on Dryad at http://dx.doi.orgXXXXX.

The summarized posterior tree resulting from the coalescent branching model deployed in BEAST 2.6.4 resolves two deeply branching lineages we delimit as *Amia calva* and *A. ocellicauda* (Fig. 1b). An annotated phylogeny with the sampling locations noted on the tips of the tree is available on Dryad at http://dx.doi.orgXXXXX. The delimitation of the two species of *Amia* shows a break in the geographic distribution along the northern Gulf of Mexico. *Amia calva* is distributed from the Pearl River in Louisiana and Mississippi, USA east including the Florida Peninsula, and the rivers draining to the Atlantic Ocean in Georgia, South Carolina, North Carolina, and Virginia, USA (Fig. 1a). *Amia ocellicauda* was first described in 1836 [S14] and is distributed from the Lake Pontchartrain system west in Gulf of Mexico draining rivers to the Colorado River system in Texas, USA, throughout the Mississippi River Basin, the Great Lakes Basin, the St. Lawrence River system, including Lake Champlain, and the Atlantic draining Connecticut River system (Fig. 1a,b).

We performed a population structure analysis to assess relative genomic ancestry with a sparse non-negative matrix factorization algorithm using the ‘snmf’ function implemented in the R package LEA v.3.0.0 [S15]. During the snmf analysis, we searched for the optimal number of ancestral populations (K) based on the cross-entropy criterion [S16], through a wide range (K = 1–20) with 10 replications for each scenario on a genotype data matrix for only biallelic, unlinked SNPs. With the R package Hierfstat [S17], we estimated the fixation index (*Fst*) for all pairs of specimens among the 177 sampled individuals.

Patterns of genomic ancestry estimated in the snmf analysis demonstrates genetic distinctiveness between the two species (Fig. 1b). Populations with signatures of admixture where those that are reconstructed as early branching in the coalescent model-inferred phylogenomic tree (Fig. 1b), which we interpret as genomic ancestral polymorphism resulting from incomplete lineage sorting. The mean *Fst* among all intraspecific comparisons of *Amia calva* and *A. ocellicauda* were less than 0.25 and the average *Fst* value among all comparisons of *A. calva* and *A. ocellicauda* was greater than 0.55 (Fig. 1c).

### Estimation of divergence times among living species of *Amia*

The divergence time of the two delimited species of *Amia* was estimated using a fossil tip dating strategy and the fossilized birth-death (FBD) branching model in BEAST 2.6.4 [S13, S18]. A total of 699 orthologous ddRAD loci were identified for the Spotted Gar *Lepisosteus oculatus* and the two species of *Amia*. A single individual from each of the three sampled species were included in the FBD fossil tip dating analysis. All loci were linked to a single tree branching model and the HKY+G model was used. The prior settings for the FBD included an exponential distribution for the diversification rate and uniform distributions for the time of origin, sampling proportion, and turnover parameters. The chain was run for 1.0 × 108 generations and log and tree files were updated every 1.0 × 104 generations. Convergence of parameters values in the BEAST 2.6.4 MCMC were assessed by the effective sample sizes that were calculated using Tracer version 1.7. Generations sampled before convergence was attained were discarded as burn-in. The BEAST analyses were run three separate times and post burn-in generations were pooled from all three runs using LogCombiner 2.8. A maximum clade credibility tree with median node heights was constructed for the post burn-in species tree topologies using TreeAnnotator 2.6.4. The xml files used in the tip dated BEAST analyses are available on Dryad at http://dx.doi.orgXXXXX. The fossil lineages of Halecomorphi used in the tip dating analysis are listed below and their phylogenetic relationships were enforced with clade constraints that reflect relationships presented in phylogenetic analyses of living and extinct lineages Holostei using morphological characters (Grande and Bemis 1998). The xml files used in the BEAST FBD analyses and summarized posterior time tree are available on Dryad at http://dx.doi.orgXXXXX.

†*Ionoscopus cyprinoides* and †*Caturus furcatus* (152.06-150.94 Ma) are from Jurassic Solnhofen, Germany [3, 4]. This site lies within the †*Hybonoticeras hybonotum* Zone [S19], the first ammonite zone of the Tithonian. This results in an age-range estimate of 152.06 to 150.94 Ma based on the interpolated ages for Jurassic ammonite biozonation [S20]. We assign an age of 151.5 Ma for both †*Ionoscopus cyprinoides* and †*Caturus furcatus*.

†*Calamopleurus cylindricus* (113-111 Ma) is from the Romualdo Member of the Santana Formation [3], which is assigned an early Albian age based on palynological studies [S21]. The end of the early Albian is placed approximately at 111 Ma [S20], while the base of the stage is placed at 113 Ma. We assign an age of 112.0 Ma for †*Calamopleurus cylindricus*.

†*Amia pattersoni* (51.66 Ma) is from the Fossil Butte Member of the Green River Formation [3]. This deposit is placed within the Wasatchian North American Land Mammal Age. Absolute dating of a tuff in the Fossil Butte Member provides an age of 51.66 ± 0.09 Ma [S22]. We assign an age of 51.66 Ma for †*A. pattersoni*.

†*Amia scutata* (35 Ma) is from the Florissant Formation, Colorado, USA and is dated at 35 million years [3]. We assign an age of 35 Ma for †*A. scutata*.

### Assessment of disparity between *Amia calva* and *A. ocellicauda* in meristic traits

To investigate if disparity in meristic traits used to discover, delimit, and describe species of fishes was consistent with the phylogenomic delimitation of *Amia calva* and *A. ocellicauda*, we collected data from 152 specimens of *Amia calva* and 73 specimens of *A. ocellicauda* following standard protocols [3, S23]. Meristic traits were the number of lateral line scales, the number of scales above the lateral line, the number of scales below lateral line at the pelvic fin, the number of transverse scale rows at the pelvic fin, the number of scale rows below the lateral line at the anal fin, the number of transverse scale rows at the anal fin, the number of scales across the breast between the pectoral fins; the number of rays, each, in the dorsal, anal, pectoral, pelvic, and caudal fins; and the number of branchiostegal rays. The meristic data for each specimen and museum collection information for each specimens are available on Dryad at http://dx.doi.orgXXXXX.

A principal components (PC) analysis of the meristic traits performed using the “prcomp” function in R version 3.2.0 (http://www.R-project.org/) showed substantial overlap of the two species when plotting PC2 vs PC1 (Fig. S2). There is no apparent geographic pattern within either of the two species. A cross-validation linear discriminant analysis (LDA) conducted with the R package MASS (https://cran.r-project.org/web/packages/MASS/index.html) shows that 88.9% of all *Amia calva* specimens are correctly identified. In contrast, only 32.2% of the specimens of *A. ocellicauda* are correctly identified using the meristic trait data.

### Characterization of morphological differences in the skulls of *Amia calva* and *A. ocellicauda*

To further assess the presence of morphological differences between *Amia calva* and *A. ocellicauda*, we scanned 8 specimens of *Amia calva* and 10 specimens of *A. ocellicauda* using high resolution computed tomography with a Nikon XT H 225 ST system. All scan parameters are provided in Table S1. Volume rendering was performed in VGStudio Max 3.5.1. We used the ImageJ software to take digital measurements of CT scans digitally rendered in VGStudio MAX 3.5. Measurements were taken of the maximum depth and length of the suborbital and interopercle, as well as of the number of alveoli in the dentary tooth row. All plots were made using ggplot2 in Rstudio. We found that interopercle robusticity (Fig. 1d) and the number of dentary teeth were consistently different in the two living species of *Amia*: *A. calva* is characterized by an elongated interopercle and 16 or 17 dentary teeth, whereas *A. ocellicauda* possesses 15 dentary teeth and a robust interopercle (Fig. 1d).

## References

1. Lidgard, S., and Love, A.C. Rethinking living fossils BioScience 68 (2018), pp. 760–770.

2. Braasch, I., Gehrke, A.R., Smith, J.J., Kawasaki, K., Manousaki, T., Pasquier, J., Amores, A., Desvignes, T., Batzel, P., Catchen, J., et al. The spotted gar genome illuminates vertebrate evolution and facilitates human-teleost comparisons Nat Genet 48 (2016), pp. 427–437.

3. Grande, L., and Bemis, W.E. A comprehensive phylogenetic study of amiid fishes (Amiidae) based on comparative skeletal anatomy. An empirical search for interconnected patterns of natural history J. Vert. Paleo. 18 (Memoir 4) (1998), pp. 1–690.

4. Grande, L. An empirical and synthetic pattern study of gars (Lepisosteiformes) and closely related species, based mostly on skeletal anatomy. The resurrection of Holostei Amer. Soc. Ich. Herp. Spec. Pub. 6 (2010), pp. 1–871.

5. Thompson, A.W., Hawkins, M.B., Parey, E., Wcisel, D.J., Ota, T., Kawasaki, K., Funk, E., Losilla, M., Fitch, O.E., Pan, Q., et al. The bowfin genome illuminates the developmental evolution of ray-finned fishes Nat Genet 53 (2021), pp. 1373–1384.

6. Sinopoli, D.A., and Stewart, D.J. A synthesis of management regulations for Bowfin, and conservation implications of a developing caviar fishery Fisheries 46 (2021), pp. 40–43.

7. Rypel, A.L., Saffarinia, P., Vaughn, C.C., Nesper, L., O’Reilly, K., Parisek, C.A., Miller, M.L., Moyle, P.B., Fangue, N.A., Bell-Tilcock, M., et al. Goodbye to “rough rish”: paradigm shift in the conservation of native fishes Fisheries 46 (2021), pp. 605–616.

8. Soltis, D.E., Morris, A.B., McLachlan, J.S., Manos, P.S., and Soltis, P.S. Comparative phylogeography of unglaciated eastern North America Mol. Ecol. 15 (2006), pp. 4261–4293.

9. Nagalingum, N.S., Marshall, C.R., Quental, T.B., Rai, H.S., Little, D.P., and Mathews, S. Recent synchronous radiation of a living fossil Science 334 (2011), pp. 796–799.

10. Kadarusman Sugeha, H.Y., Pouyaud, L., Hocdé, R., Hismayasari, I.B., Gunaisah, E., Widiarto, S.B., Arafat, G., Widyasari, F., Mouillot, D., et al. A thirteen-million-year divergence between two lineages of Indonesian coelacanths Sci Rep 10 (2020), p. 192.

## References

S11. Peterson, B.K., Weber, J.N., Kay, E.H., Fisher, H.S., and Hoekstra, H.E. Double digest RADseq: an inexpensive method for ee novo SNP discovery and genotyping in model and non-model species Plos One 7(5) (2012), p. e37135.

S12. Eaton, D.A.R., and Overcast, I. ipyrad: Interactive assembly and analysis of RADseq datasets Bioinformatics 36 (2020), pp. 2592–2594.

S13. Bouckaert, R., Vaughan, T.G., Barido-Sottani, J., Duchêne, S., Fourment, M., Gavryushkina, A., Heled, J., Jones, G., Kühnert, D., De Maio, N., et al. BEAST 2.5: An advanced software platform for Bayesian evolutionary analysis Plos Comput Biol 15 (2019), p. e1006650.

S14. Richardson, J. (1836). Fauna Boreali-Americana; or the zoology of the northern parts of British America: containing descriptions of the objects of natural history collected on the late northern land expeditions, under the command of Sir John Franklin, R. N. Part 3, (London: J. Bentley).

S15. Frichot, E., and François, O. LEA: An R package for landscape and ecological association studies Methods Ecol Evol 6 (2015), pp. 925–929.

S16. Alexander, D.H., and Lange, K. Enhancements to the ADMIXTURE algorithm for individual ancestry estimation BMC Bioinf. 12 (2011), p. 246.

S17. Goudet, J. hierfstat, a package for R to compute and test hierarchical F-statistics Mol Ecol Notes 5 (2005), pp. 184–186.

S18. Gavryushkina, A., Heath, T.A., Ksepka, D.T., Stadler, T., Welch, D., and Drummond, A.J. Bayesian total-evidence dating reveals the recent crown radiation of penguins Syst. Biol. 66 (2017), pp. 57–73.

S19. Schweigert, G. Ammonite biostratigraphy as a tool for dating Upper Jurassic lithographic limestones from South Germany first results and open questions Neues Jahrbuch für Geologie und Paläontologie - Abhandlungen 245 (2007), pp. 117–125.

S20. Gradstein, F.M., Agterberg, F.P., Ogg, J.G., Hardenbol, J., Veen, P.V., Thierry, J., and Huang, Z. (1995). A Triassic, Jurassic, and Cretaceous time scale. In Geocrhonology, time scales and global stratigraphic correlation, Volume 54, W.A. Berggren, D.V. Kent, M.-P. Aubry and J. Hardenbol, eds. (Tulsa, Oklahoma: SEPM), pp. 95–126.

S21. Heimhofer, U., and Hochuli, P.A. Early Cretaceous angiosperm pollen from a low-latitude succession (Araripe Basin, NE Brazil) Rev. Palaeobot. Palyno. 161 (2010), pp. 105–126.

S22. Smith, M.E., Carroll, A.R., and Singer, B.S. Synoptic reconstruction of a major ancient lake system: Eocene Green River Formation, western United States Geol. Soc. Am. Bull. 120 (2008), pp. 54–84.

S23. Hubbs, C.L., Lagler, K.F., and Smith, G.R. (2004). Fishes of the Great Lakes Region, (Ann Arbor: The University of Michigan).

